# Genomes of the Golden Horde Elites and their Implications for the Rulers of the Mongol Empire

**DOI:** 10.1101/2025.06.11.659160

**Authors:** Ayken Askapuli, Hideaki Kanzawa-Kiriyama, Tsuneo Kakuda, Aibar Kassenali, Syrym Yessen, Uli Schamiloglu, Steven J. Schrodi, John Hawks, Naruya Saitou

**Affiliations:** Department of Integrative Biology, University of Wisconsin-Madison, Wisconsin, United States of America; Department of Anthropology, National Museum of Nature and Science, Tsukuba, Japan; Center for the Study and Preservation of Cultural Heritage, Astana, Kazakstan; Astana International University, Astana, Kazakstan; Department of Legal Medicine, University of Yamanashi, Chuo City, Japan; School of Sciences and Humanities, Nazarbayev University, Astana, Kazakstan; Department of Medical Genetics, University of Wisconsin-Madison, Wisconsin, United States of America; Department of Anthropology, University of Wisconsin-Madison, Wisconsin, United States of America; National Institute of Genetics, Mishima, Japan

## Abstract

The Golden Horde, the northwestern extension of the Mongol Empire that was ruled by the eldest son of Genghis Khan (also spelled Chinggis Khan) and his descendants, played an important role in the history of Central Eurasia and eastern Europe. For this reason, the genetic profiles of Genghis Khan and his descendants are of keen academic and public interest. His Y-chromosome has been hypothesized to belong to haplogroup C3* (C2a1a3-F1918) but this has yet to be substantiated. To explore the genetic ancestry of Genghis Khan, we analyzed four archaeological individuals from the Golden Horde, including three males and one female, who were buried in separate medieval elite mausoleums in the Ulitau region of Kazakstan. One of the males is believed to be Joshi (Juchi), the eldest son of Genghis Khan. We generated genomic data from these archaeological individuals and found that the three males were paternally related, sharing the Y-chromosome haplogroup C3*, consistent with the hypothesized signature haplotype of Genghis Khan. In addition, we confirm that the Golden Horde elites derived most of their genomes from Ancient Northeast Asians (ANA) while having an additional ancestral component from either Ancient North Eurasians (ANE) or a Berel Scythian related population. Furthermore, we were able to identify their medieval relatives in the Mongolian Plateau through constructing an Identity by Descent (IBD) network.

## Introduction

The Mongol Empire developed by Genghis Khan and his descendants had a profound impact on the political and economic order of Eurasia. In the peaceful period termed as *the Pax Mongolica* following its conquest, the Mongol Empire contributed significantly to the development of political institutions, economic relations, and cultural diversity of the region^1^, thereby facilitating transregional relations and trade between Asia and Europe^2,3^.

The Golden Horde, the northwestern extension of the Mongol Empire, stretched from the Ertis River (Irtysh) in the east to the Pontic-Caspian steppe in the west^4^, with its center of gravity located in the west^1^. While the Khanate has been known to the West as the *“Golden Horde”*, it was also known as the *Ulug Ulus*, *Kipchak Khanate*, or *Ulus Juchi.* The capital of Golden Horde was first Saray Batu, then Saray Berke, both situated on the bank of the Edil River (now Volga)^4^. The initial orda (residence) of Joshi Khan that later became the capital of the White Horde, a subdivision of the Golden Horde, was located on the bank of the Ertis River, as recorded by John de Plano Carpini in the 13th century^5^. The Golden Horde ruled the Kipchak steppe, western Siberia, and the Pontic-Caspian region from 1240 to 1480 CE, which lasted one century longer than the Mongol rule in China and Persia. The Mongol Empire and its successor states continued their long existence as a result of the administrative flexibility of the nomads and the legacy of the political acumen of Genghis Khan^1^.

The historical importance of the Golden Horde in Eurasia has been well appreciated. The Golden Horde created a symbiosis of nomadic and sedentary cultures^1^, enabling ethnically and culturally diverse peoples with distinct confessions to live peacefully in its domain, thanks to the religious tolerance of the Golden Horde rulers. Moreover, the impact of the Golden Horde on the Kievan Rus should not be underestimated, as evidenced by the impact of the Golden Horde on the economy, political system, and culture of the eastern Slavs^1^.

Along with its geo-political and commercial influences, the Golden Horde played a critical role in the emergence of modern ethno-linguistic groups in Central Eurasia, including Tatars, Nogays, Özbeks, Kazaks, and others. The ethno-linguistic identities of these groups were formed and further consolidated in the Khanates that emerged out of the disintegration of the Golden Horde. The immediate successor states included the Kazan Khanate, the Crimean Khanate, the Astrakhan Khanate, the Shibanid Khanate in Siberia, the Muscovite client state of Kasimov, the nomadic Nogay Horde, and the Great Horde, which was the nomadic core of the Golden Horde^1^. It is well known from historical annals that, with few exceptions, the ruling elites of the Golden Horde and its successor states trace their ancestry to Joshi, the oldest son of Genghis Khan^1^. Furthermore, some of the ethnonyms, such as Nogay and Özbek, are associated with the personal names of the Golden Horde leaders.

The Ulitau massif in central Kazakstan boasts many historical relics from the Golden Horde period, including numerous burial complexes and impressive medieval mausoleums, which bear witness to the former historical importance of the region. One of the mausoleums is named today after Joshi. Kazaks often pay tribute to the mausoleum, believing that the person interred in the mausoleum was the eldest son of Genghis Khan. However, some scholars are skeptical of this claim, arguing that, even though the mausoleum was named after Joshi, the man buried inside might not be Joshi himself^6,7^. As an example, the historian Bartold argued that the mausoleum was dedicated to Joshi posthumously^6^.

We know from historical annals that Joshi died in 1227 CE, several months before his father, Genghis Khan, passed away^8,9^. According to the later historical tradition of the Golden Horde, Joshi Khan died when he fell from a horse at Ulitau (Ұлытау, Ulutau, Uligh Tagh), where he was then buried^10^. According to Kazak folklore, Joshi Khan died of an accident while chasing wild horses, and the musical composition *<u>Aksak Kulan</u>* (literally *lame wild horse)* was dedicated to this incident^11^.

Despite the historical importance of Joshi and his descendants, their genetic origins as well as that of Genghis Khan remained mysterious. Historical records provide us with contradictory information about the ancestral origin of Genghis Khan. His portrait, painted during the Mongol Empire and now stored in the National Palace Museum of Taiwan^12^, shows typical Mongolian traits with prominent cheekbones, a flat face, almond-shaped eyes, a broad forehead, and a scarce beard. By contrast, the historian Rashid al-Din describes Genghis Khan as having blue eyes and a heavy beard^13^, which evokes a more European-looking image of Genghis Khan.

One seemingly unsurmountable hurdle in revealing his genetic origins is the difficulty of locating the archaeological remains of Genghis Khan and any of his offspring. There have been international efforts to find the burial place of Genghis Khan^14–16^, but the exact location of his tomb still eludes researchers. Historical records inform us that Genghis Khan and many of his descendants, including his grandson, Kubilay Khan, were buried in a secret site in Mongolia^6^. Instead of erecting any monuments or mausoleums, the elites were placed into a burial pit, and the burial earth was flattened by the hooves of horses, after which grass seeds were sown to grow there. As a result, the burial site would blend in with surrounding grass patches, resembling the intact grassland^6^.

A previous study of Y-chromosome diversity in human populations in Asia hypothesized that Genghis Khan was responsible for the high frequency of haplogroup C3 Star Cluster (C3*)^17^. However, a later ancient DNA study performed on elite burials of the Mongol Empire from Sukhbaatar, eastern Mongolia, suggested Y-chromosome haplogroup R1b was the paternal lineage of Genghis Khan^18^. Yet, another ancient DNA study identified Y-chromosome haplogroup Q in archaeological individuals interred in a Mongolian noble family mausoleum in Hebei, northern China^19^. A more recent ancient DNA study by Jeong and colleagues^20^ reported genomic data from more than 200 individuals who lived in the Mongolian Plateau from the pre-Bronze Age through the medieval periods, and revealed a heterogeneous prehistoric genetic landscape of that region, implying that Rashid al-Din’s description of Genghis Khan cannot be simply dismissed as baseless.

In this study, we aimed to disentangle the genetic origins of the ruling elite of the Mongol Empire and evaluate the folklore-based hypothesis that the archaeological individuals in Ulitau were of Mongolian origin, and thus trace their ancestry to the Mongolian Plateau. An alternative hypothesis is that these individuals have local ancestry, akin to the medieval Kipchaks. Here, we report genomic data from four archaeological individuals recovered from Ulitau — three males and one female — who most likely belonged to the elite class of the Golden Horde, as evidenced by the spectacular mausoleums dedicated to them. The genomic ancestry of these individuals shows that they had genetic affinity with medieval individuals from the Mongolian Plateau, clearly indicating their Mongolian origin. Moreover, the three males harbored the identical paternal lineage, C3*, that was previously hypothesized to be the paternal lineage of Genghis Khan^17^.

## Results

### Archaeological artifacts

Archaeological individuals were excavated from four medieval mausoleums in the Ulitau massif, central Kazakstan (Fig 1a). The architectural styles of these mausoleums were well described during the Soviet period^21^. The burials in the mausoleums of Joshi and Alasha Khan were excavated in the Soviet era (Fig S1) and no artifacts of special interest or value were reported^22^. The archaeological findings were not published but survived as a report^23^. According to this report, one male was buried in the Mausoleum of Alasha Khan, accompanied with azure glassware that was broken. Anthropologist Orazak Ismagulov analyzed the skeletal remains of Alasha Khan and his team performed craniofacial reconstruction of this individual^24^.

**Fig 1.**
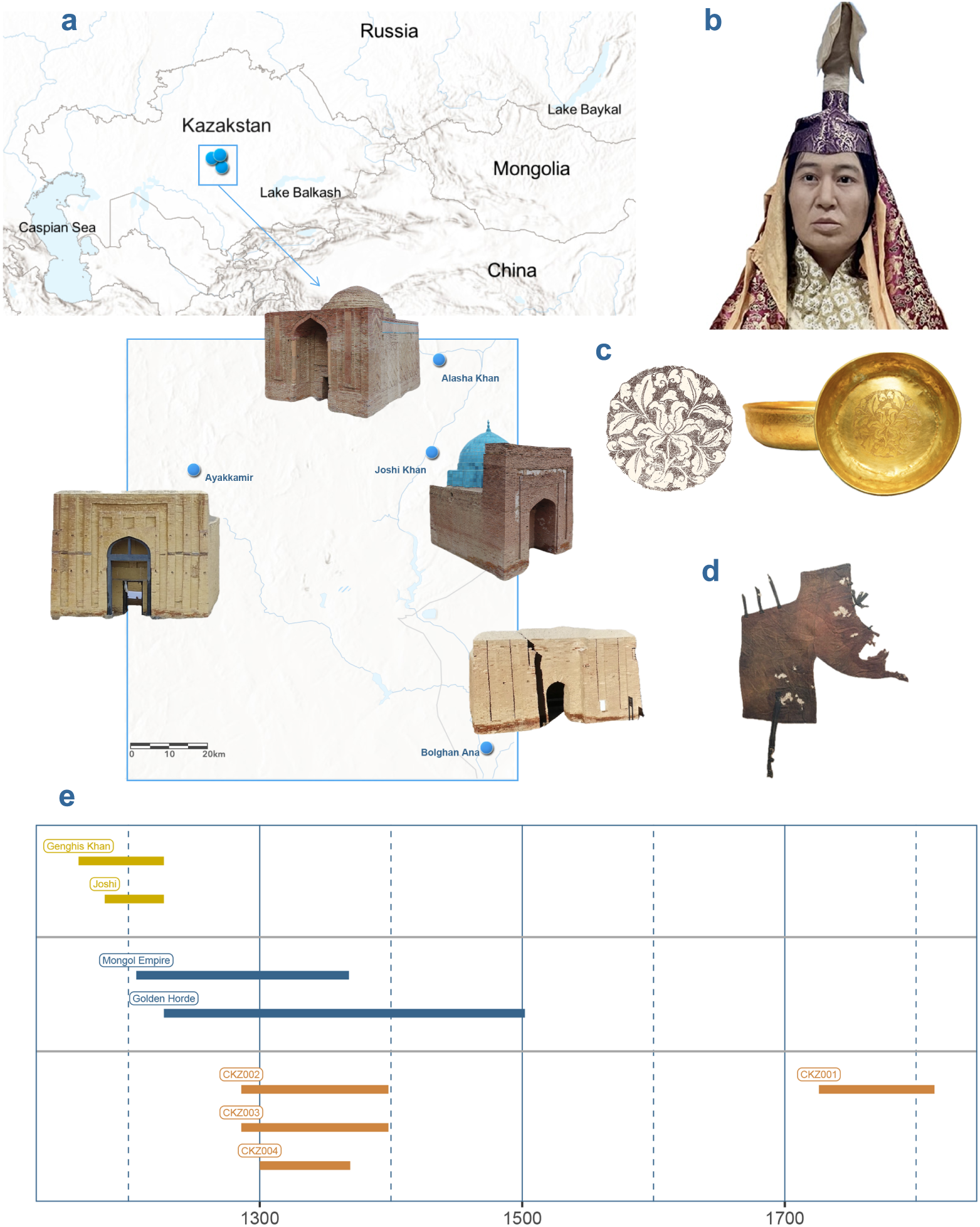
Archaeological sites and radiocarbon dates. **Notes: a.** Archaeological site map of the mausoleums; the orientations of the mausoleums are given as they were pictured. **b.** Reconstructed image of Bolghan Ana; **c.** Golden cup and its lotus motif; **d.** a cloth fragment; **e.** Radiocarbon dates contrasted with the lifespan of historical figures and polities. The collapse of the Great Horde in 1502 marks the end of the Golden Horde. CKZ001: male from the Mausoleum of Alasha Khan; CKZ002: male from the Mausoleum of Joshi; CKZ003: male from the Ayakkamir Mausoleum; and CKZ004: female from the Mausoleum of Bolghan Ana *(could also be spelled Bulghin or Bulgan Ana*^6^).

In the Mausoleum of Joshi, there were two individuals — one male and one female — buried along with a camel’s head. In addition, animal bones and fragments of clothes and leather were discovered in the tombs. Anthropologist Vulf Ginzburg analyzed the skeletal remains of the male from this Mausoleum and concluded that he might be a descendant of Joshi^25^.

The other two mausoleums, Bolghan Ana and Ayakkamir, were excavated for the first time. From Bolghan Ana, we discovered a saddle with gears, a pair of golden earings, a bone hair stick, fragments of clothes, a saber, and a golden cup (Fig S2 & S3; Fig 1c & d)^26^. Among the artifacts, the golden cup was the most informative. The inside of the cup was decorated with a lotus flower motif. The use of lotus as a decorative motif predates the Mongol Empire^27^. A golden cup with a similar lotus motif was previously unearthed in Siberia and attributed to the Golden Horde^28^.

In the Ayakkamir Mausoleum, no artifacts were discovered except for a bronze cup. The skull of the individual was missing, possibly being taken in a previous excavation, since no marks of beheading were observed in the cervical vertebrae.

One common feature of all four burials is that human remains were oriented towards west or northwest, a burial practice clearly influenced by Islam. In a medieval burial called Karasuuir (Қарасуыр, DA28 in Fig 5 & 8 from this burial site), also from Ulitau, the human corpse was oriented to the north^29^. Placing deceased individuals in the north or northeast orientation is a typical feature of the burials from the Mongol Empire^30^. Since the Mausoleum of Joshi is located on the bank of the Kenggir (Kara-Kenggir) River, we designated all four archaeological individuals as *Kenggir individuals*.

### Radiocarbon dating

We obtained radiocarbon (C^14^) dates for bone fragments from the human remains (Fig 1e and Table 1). The C^14^ dating was carried out by Beta Analytic, Miami, FL, USA. Except for CKZ001, the radiocarbon dating results agree with the period of the Golden Horde. While CKZ001 was dated to as recently as the 18^th^ century CE, the other three individuals, CKZ002, CKZ003, and CKZ004, were dated to the 14^th^ century CE, with CKZ002 and CKZ003 having the same estimates (Table 1). A recent radiocarbon study^31^ on the wooden materials used for the construction of the Mausoleum of Joshi gave a date range (1298-1398 CE) comparable to our estimates. However, the wood used for the coffin had slightly older radiocarbon age (∼1245 CE)^31^.

**Table 1.** Information on the archaeological individuals from Ulitau, Central Kazakstan.

| Sample ID | Estimated Age (95% CI) | Molecular Sex | 1240K SNPs | Paternal Lineage | Maternal Lineage | Read Depth |
| --- | --- | --- | --- | --- | --- | --- |
| CKZ001 | 1726-1814 cal AD | Male | 1149K | C2a1a3 | D4c2b | 14.5936 |
| CKZ002 | 1286-1398 cal AD | Male | 1001K | C2a1a3 | C4a2a1 | 3.88456 |
| CKZ003 | 1286-1398 cal AD | Male | 636K | C2a1a3 | C4a1a+195 | 0.793282 |
| CKZ004 | 1300-1369 cal AD | Female | 1112K | NA | G2a1g | 3.88796 |

### Ancient DNA extraction and sequencing

We were able to extract DNA from samples of CKZ001 (tooth), CKZ002 (petrous bone), CKZ003 (talus), and CKZ004 (tooth), albeit with miniscule amount of DNA from the petrous bone of CKZ002. Therefore, we performed an additional DNA extraction from a fibula fragment of CKZ002, given the importance of this individual. As such, we prepared several aDNA libraries from CKZ002 and sequenced them separately (Table S1). The combined aDNA data from the libraries gave a read depth of ∼ 3.9X, enabling us to call a sufficient number of 1240K SNPs^32^ for CKZ002 (Table 1). We obtained the highest quality aDNA data (∼ 14.6x read depth) from CKZ001, in alignment with its relatively recent radiocarbon age, and thus experiencing less DNA damage.

### Principal component analysis (PCA)

To understand the genetic affinity of the Kenggir individuals with modern and ancient human populations in Eurasia, we performed an analysis on autosomal genetic variants (Chr1–22) via smartPCA^33^, which can calculate principal components using modern individuals and projects ancient individuals on top of the PCA space.

In the PC plane of PC1 and PC2, as shown in Fig 2, the Kenggir individuals clustered together, indicating their shared genomic background. In the PCA space, the Kenggir individuals positioned themselves in proximity with modern populations from Siberia (Altaians, Buryats, Dolgans, and Yakuts), and northeast Asia (Ulch, Daur, Oroqen, etc.). Kalmaks were the only population from further west that clustered close to the Kenggir individuals. This is likely due to the fact that Kalmaks migrated to the north of the Caspian Sea from the Altay-Sayan region in the east just a few centuries ago. All of these populations are Altaic speaking ethnic groups, with Altaians, Dolgans, and Yakuts (Saka) speaking Turkic languages, the Buryats, Daur, and Kalmaks speaking Mongolic languages, and the Ulch and Oroqen (Orochen) speak Tungusic languages.

**Fig 2.**
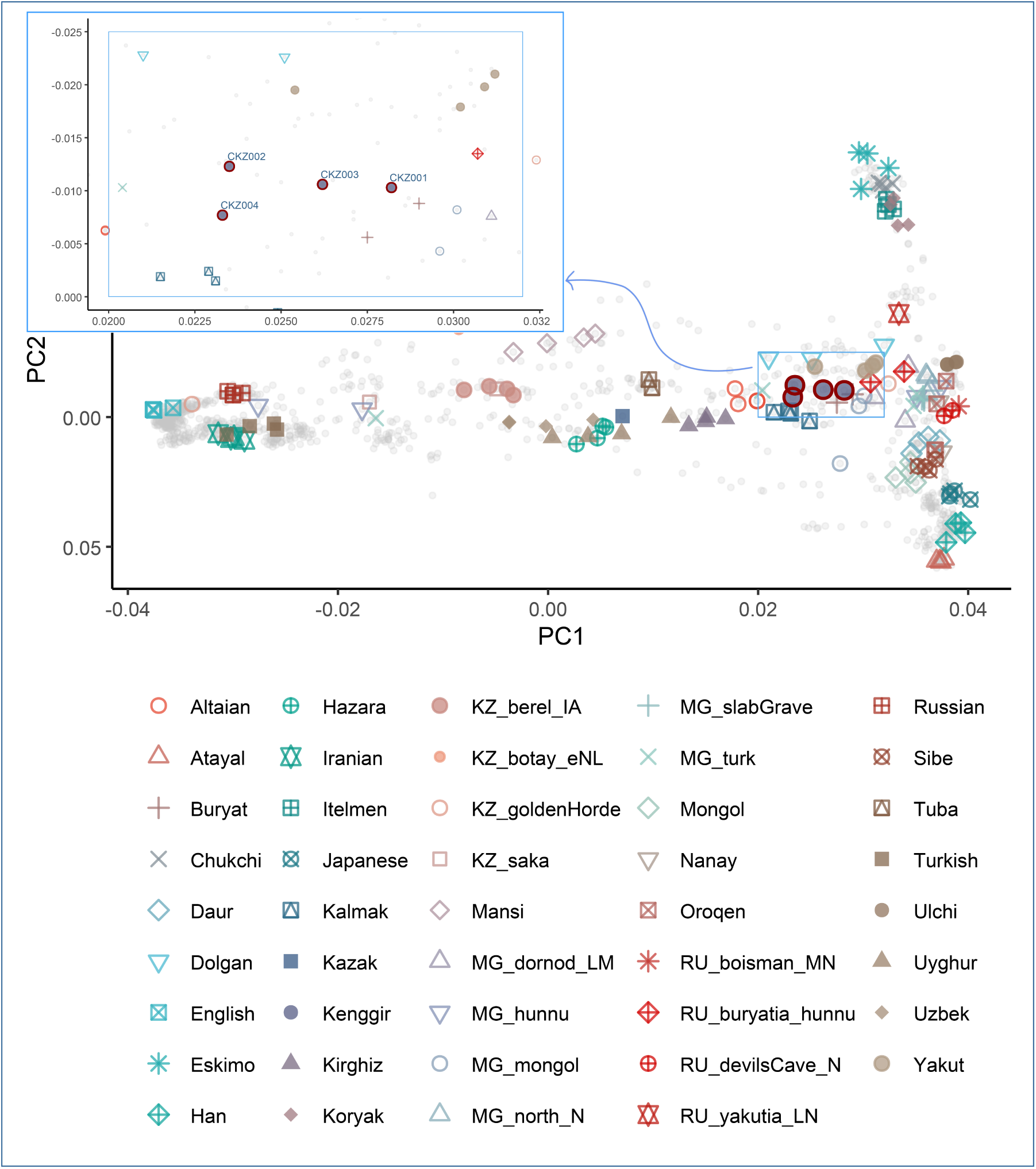
PCA analysis together with ancient and modern human populations from Central Eurasia. **Note:** The dark red circles indicate the Kenggir individuals.

A few ancient individuals from the Mongolian Plateau and the Kazak Steppe also clustered close to the Kenggir individuals in the PCA space. The analysis also showed the genetic relatedness of the Kenggir individuals and Hunnu period individuals from Buryatia, the Middle Neolithic Boisman individual, and the Neolithic individuals from the Devil’s Cave, northeast Asia (sample details in Table S2).

### *f*-statistics

Aiming to understand the genomic makeup of the four individuals from Golden Horde in the context of the Central Eurasian genomic landscape, we performed qp3Pop and qpDstat analyses^34^ and calculated *f*-statistics. The outgroup *f_3_*-statistic (Fig 3) and *f_4_*-statistic (Fig 4) indicated that the Kenggir individuals were much closely related with populations from northeast Asia (Ulch, Oroqen, Nanay, and Daur) and northern Siberia (Yakut and Even), in agreement with the PCA results.

**Fig 3.**
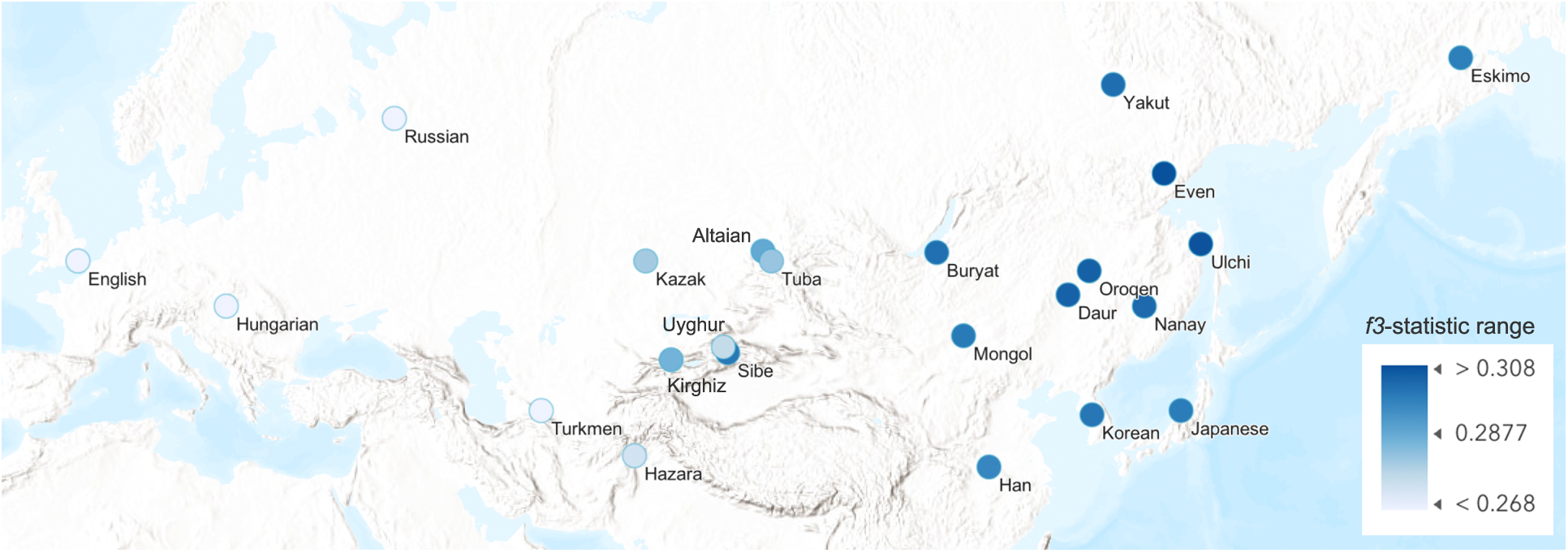
Outgroup *f3*-statistic calculated with the model (CKZ002, X; Mbuti) **Notes:** Only modern individuals were included in the plot. The darker blue color indicates closer genetic affinity between the population and the Kenggir individuals, represented by CKZ002 in this figure.

**Fig 4.**
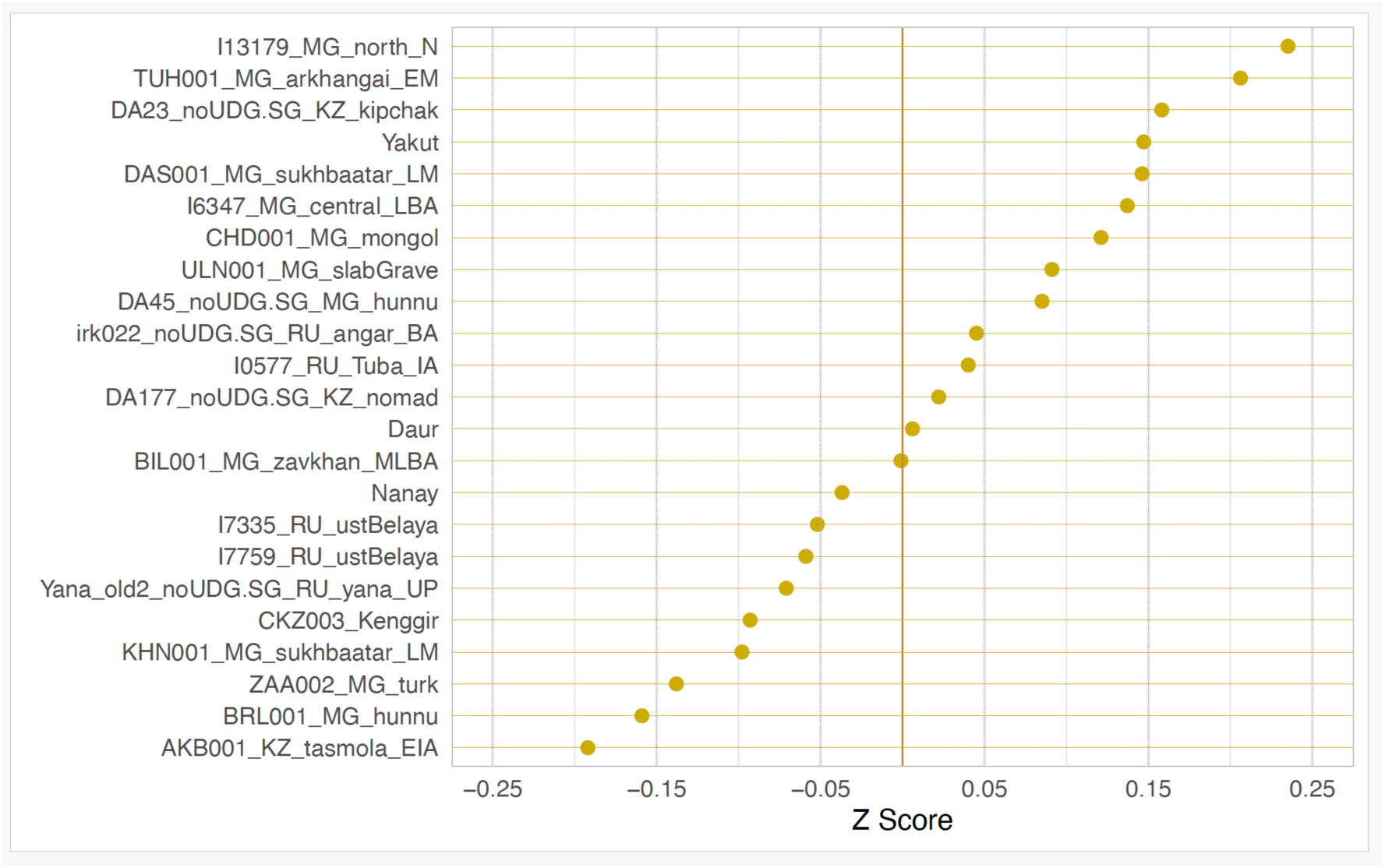
Z-scores of the *f4*-statistic calculated from the model (Mbuti, Ust-Ishim; X, CKZ002)

**Fig 5.**
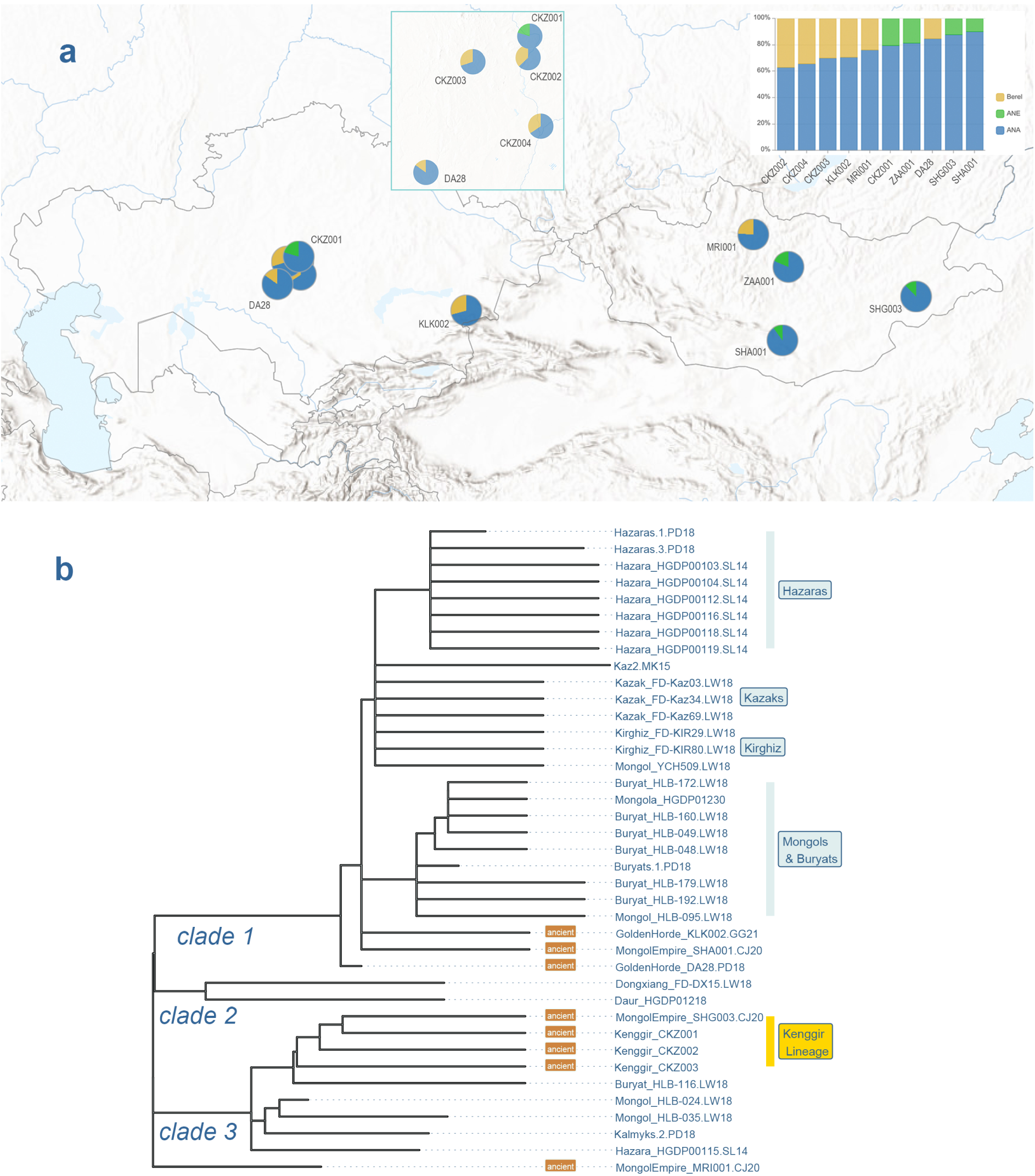
Modeling genetic ancestry based on autosomes and Y-chromosome. **Notes**: Clades 1 and 2 are the two sub-branches of C2a1a3a1-F5481, while Clade 3 corresponds to C2a1a3a6-SK1072. Y-chromosome sequences are from PD18: Damgaard et al. 2018^40^; SL14: Lippold et al. 2014^41^; LW18: Wei et al. 2018^42^; GG21: Gnecchi-Ruscone et al. 2021^43^; and CJ20: Jeong et al. 2020^20^.

In the outgroup *f_3_* statistic, the population pairs with similar genetic drift from the reference population (i.e., Mbuti, a hunter-gatherer group, from the Congo region of Africa) showed the highest *f_3_* values, as indicated by the darkest circles in Fig 3. For the *f_4_*-statistic (Fig 4), the Altaic speaking modern populations (e.g., Daur, Nanay, and Yakut) exhibited *f_4_* values and corresponding Z-scores that did not significantly deviate from zero, indicating a possible clade relationship with the Kenggir individuals. The genetic affinity of the Daurs with the Kenggir individuals was also reflected in their language, which has retained more medieval Mongolian linguistic features^8^. In addition, for the *f_4_*-statistics, ancient individuals from Mongolia who belonged to various archaeological cultures, including Hunnu, Slab Grave, Türük, and the medieval period, showed close genetic affinity with the Kenggir individuals. This result indicated genetic continuity of some populations across various archaeological ages, despite the overall genetic heterogeneity observed in the Mongolian Plateau.^20^

### Modeling the ancestry of ruling elites of the Golden Horde

After identifying the populations and individuals who showed genetic affinity with the Kenggir individuals through PCA and outgroup *f_3_*-statistic, we wanted to identify the ancestral populations that made significant contributions to the gene pool of the Kenggir individuals. To this end, we performed modeling with qpAdm^34,35^. For comparability of our results with those of Jeong and colleagues^20^, we used the same set of reference populations but added an additional reference, Ust-Ishim^36^. The full list of reference populations is given in the Methods section.

First, we identified ancient individuals who shared ancestry and formed a clade with CKZ002. qpAdm analysis revealed that CKZ002 formed a clade only with CKZ004, as reflected in the PCA, while the other two individuals, CKZ001 and CKZ003, did not do so (Table S4). In addition, CKZ002 also showed clade level genetic affinity with individuals from medieval Mongolia: MRI001 and ZAA001^20^. A shared feature of Kenggir individuals is that all of them derived most of their genetic makeup from ANA (Ancient Northeast Asians)^37^. The second population that contributed to the gene pool of the Kenggir individuals was a population that resemble the Berel Scythians, i.e., all but CKZ001 can be modeled to derive their ancestry from ANA and Berel. By contrast, CKZ001 received a second genetic component from ANE^38^ (Ancient North Eurasians, Fig 5a). Our analysis indicated ANE as the second genetic component of CKZ001, ZAA001, SHA001, and SHG003, although the latter three individuals were not inferred to have ANE ancestry when they were previously analyzed^20^. Our results are not unexpected given that when different combinations of reference and source populations are used, qpAdm can provide equally likely models about ancestral genetic components^35^.

### Y-chromosome haplotypes

Identification of the paternal lineage of the male from the Mausoleum of Joshi was of critical importance to us as it would shed light on the identity of the individuals as well as test the C3* haplotype hypothesis^17^.

The Identification of Y-chromosome haplotypes and reconstruction of phylogenetic tree was performed with pathPhynder^39^. The program places ancient Y-chromosomes, which usually have high missingness for SNP data, on a Y-chromosome phylogenetic tree that is built with high coverage Y-chromosome sequences from modern individuals.

The analysis revealed that the three Kenggir males (CKZ001, CKZ002, and CKZ003) were paternally related and shared a recent paternal ancestor who harbored a C3* (C2a1a3-F1918) Y-chromosome that is believed to be associated with Genghis Khan^17^. For comparative purposes, we included C3* Y-chromosomes from both modern and ancient individuals in the analysis and searched for individuals who had paternal genetic affinity with the Kenggir individuals.

However, we lacked Y-chromosome sequences from members of the Töre clan of Kazaks^44^, i.e., the golden lineage (Altan Urug) of Kazaks. By genealogy, the Töre clan members trace their origin to Joshi, the eldest son of Genghis Khan. A previous study reported that the haplotype C2a1a1b-F1756 characterizes the Töre clan^44^.

The placement of ancient Y-chromosomes in the tree (Fig 5b) was quite robust, given that hundreds of SNPs supported that branching pattern (Table S3). However, the relative positions of the external nodes could be prone to errors, since ancient samples had various levels of missingness. For instance, CKZ003 was positioned as a basal branch relative to CKZ001 and CKZ002. This might not be correct given the fact that aDNA from CKZ003 had the lowest sequence depth (Table 1), and thus only about 400 Y-chromosome markers supported this branch whereas about 1000 markers were in favor of the branching of CKZ001 and CKZ002 (Table S3).

Kenggir individuals had varying degrees of paternal relatedness with ancient individuals who lived in the same archeological timeframe. DA28, excavated from the Ulitau region^40^, harbored the same haplogroup but formed a clade with KLK002, which was excavated from a mausoleum near the medieval town of Koylik (aka Kayalyk)^43^, where Carpini sojourned in the 13^th^ century^5^. The medieval Mongolian SHA001 and Golden Horde individuals DA28 and KLK002, together with the modern Mongolic speaking Buryats and Mongols, Turkic speaking Kazaks and Kirghiz, and the Persian speaking Hazara, formed clade 1 in the Y-chromosome tree (Fig 5b). The Kazak individuals included in the analysis were from the Kerey clan^42^, which had the highest frequency of C3* Y-chromosomes^45^. The Y-chromosome haplotype was found in other Kazak clans, including Jalayirs, Dulats, and others^46–48^. Previous studies reported that C3* Y-chromosomes were found in Hazaras as well^17^.

In the phylogenetic tree, SHG003 from eastern Mongolia^20^ formed a clade with the Kenggir individuals, indicating that SHG003 was paternally closely related to them. We found two additional males from medieval Mongolia with C3* Y-chromosomes, indicating wide geographic dispersal of the lineage.

Like all of the samples in the C3* tree (Fig 5b), Kenggir males had the derived allele T for the marker F3796, which defines C3*^42^, along with upstream F1918 SNP and other markers. We define the *Kenggir lineage* as C2a1a3a6-ZQ369, based on the most recent ISOGG Y-chromosome tree (v15.73)^49^. C2-F3796 has previously been found at high frequencies in Kazak clans such as Kerey, Shapirashti, Jalayir, and others^46–48^. As expected, the Kenggir males also shared paternal ancestry with modern Buryat, Kalmak, and Mongol individuals, as shown in the clade 3 (Fig 5b). The coalescence time of C3* (C2-F3796) was previously estimated to be ∼2,500 years old^42^, apparently predating the Mongol Empire.

### mtDNA haplotypes

The four individuals were not closely related through their maternal lineages. One individual, CKZ001, had D4 mtDNA lineage, and two individuals (CKZ002 and CKZ003) belonged to subbranches of haplogroup C4, while the female (CKZ004) had G2a haplogroup (Table 1; Fig 6).

**Fig 6.**
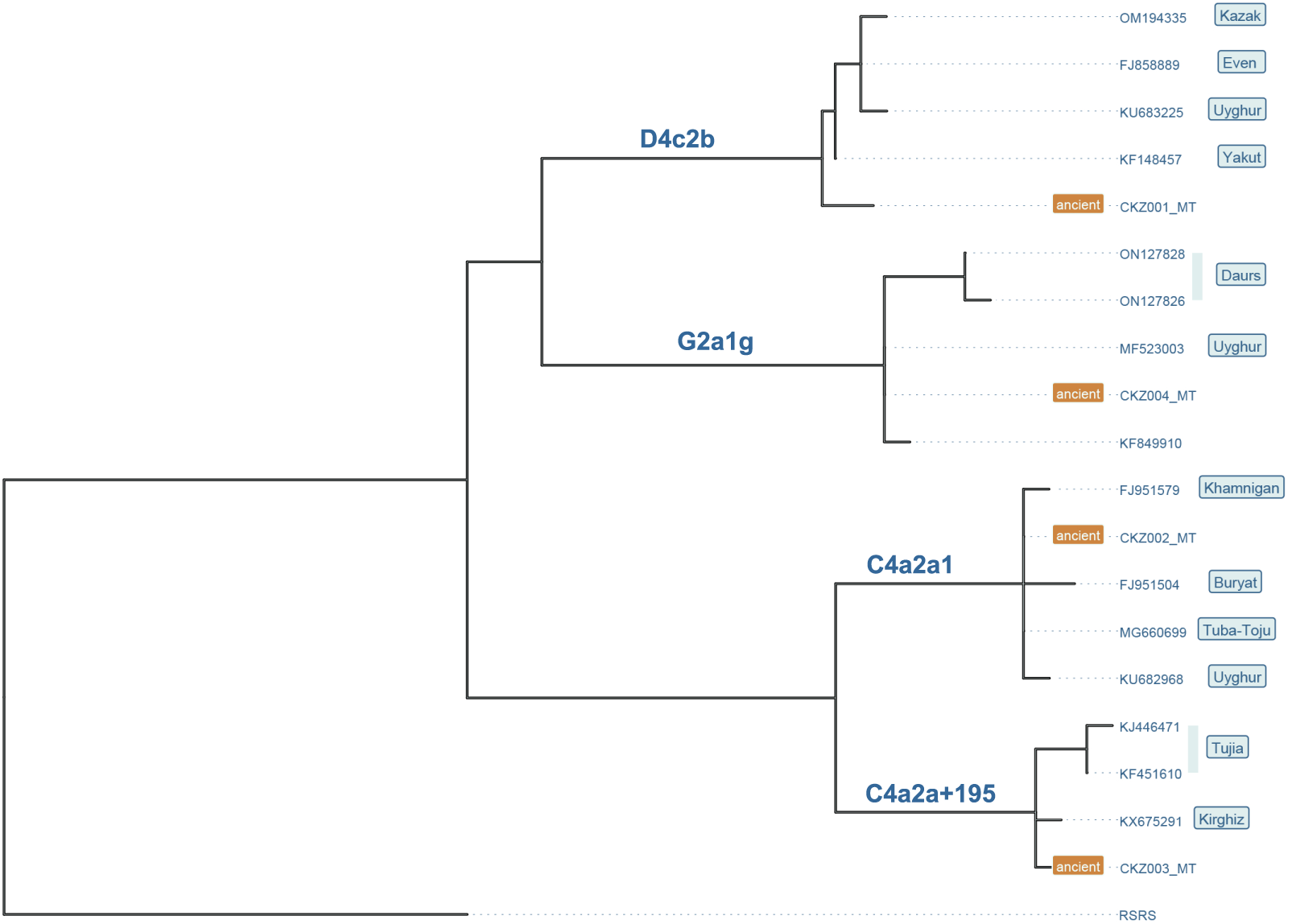
Maternal lineages found in Kenggir individuals. **Notes:** RSRS (Reconstructed Sapiens Reference Sequence)^50^ was used as an outgroup for the tree reconstruction. The ethnic origin of the sequence KF849910 was unknown.

D4 is a common East Eurasian haplogroup found at high frequencies in Kazaks and other Turko-Mongolian speakers^51–55^. The East Eurasian lineages C4 and G2 occur at relatively high frequencies in modern Turko-Mongolian populations, as well^51–53,56^.

We performed a BLAST search on GenBank and retrieved mtGenomes that had the highest levels of sequence similarity with those of the Kenggir individuals. The phylogenetic tree (Fig 6) shows the maternal genetic affinity between the Kenggir individuals and modern individuals of various ethnic origins. However, all of these individuals belong to Altaic speaking ethnic groups, including Mongolic speaking Buryats, Daurs, and Tujia; Tungusic speaking Evens; and Turkic speaking Kazaks, Kirghgiz, Tuba-Toju, Uyghurs, and Yakuts. Khamingan is a bilingual ethnic group from the east of Lake Baikal whose members speak Mongolic and Tungusic.

mtDNA heterogeneity in this region is expected, given the fact that nomadic groups in Central Eurasia practiced and still practice exogamy. Modern Kazaks and other nomadic groups in Central Eurasia exhibit high levels of diversity in mtDNA lineages, while having relatively homogenous Y-chromosome lineages^52,57^. Exogamy likely have evolved early in the nomadic groups and continued into the recent past, as evidenced by the genetic profiles of the Avars ^58^, an early medieval nomadic group.^58^

### Kinship among the Kenggir individuals

We wanted to understand the possible kinship among the three males and their genetic relationship with the female, *Bolghan Ana.* We carried out a kinship analysis using the READ program^59^ and ancIBD. As shown in Fig S4 and S5, the four individuals did not have first- or second-degree relationship, i.e., they did not have father-child, sibling, or grandparent-grandchild relationships.

According to Kazak folklore, KZ004 ( Bolghan Ana) is believed to be the daughter of CKZ002 (Joshi Khan). However, the whole genome-based kinship analyses (READ and ancIBD) did not support this claim. ancIBD analysis revealed that CKZ003 was closely related to CKZ001 (Alasha Khan) and CKZ002 (Joshi Khan), but CKZ001 and CKZ002 were relatively more distantly related, sharing only one ∼ 9 cM IBD segment (Fig S5 and S6). CKZ001 and CKZ003 shared six IBD segments totaling 121.6 cM. Based on the pattern of the IBD sharing (Fig 7), the Y-chromosome tree (Fig 5b), and the radiocarbon dates (Fig 1e and Table 1), it could be inferred that CKZ003 is ancestral to CKZ001, with approximately 4–5 generations (100–150 years) separating them, or about half of the timespan estimated by C^14^ analysis (Fig 1e). Based on IBD information, it is not possible to identify the exact relationship of pairs beyond 3^rd^ degree relatives^60^. Even though qpAdm analysis indicated a possible clade relationship between CKZ002 and CKZ004, they did not share any detectable IBD segments. The clade relationship revealed by qpAdm might be due to the shared genomic background of the individuals (Fig 5a), with CKZ002 and CKZ004 having comparable amounts of ANA and Berel Scythian related ancestral components.

**Fig 7.**
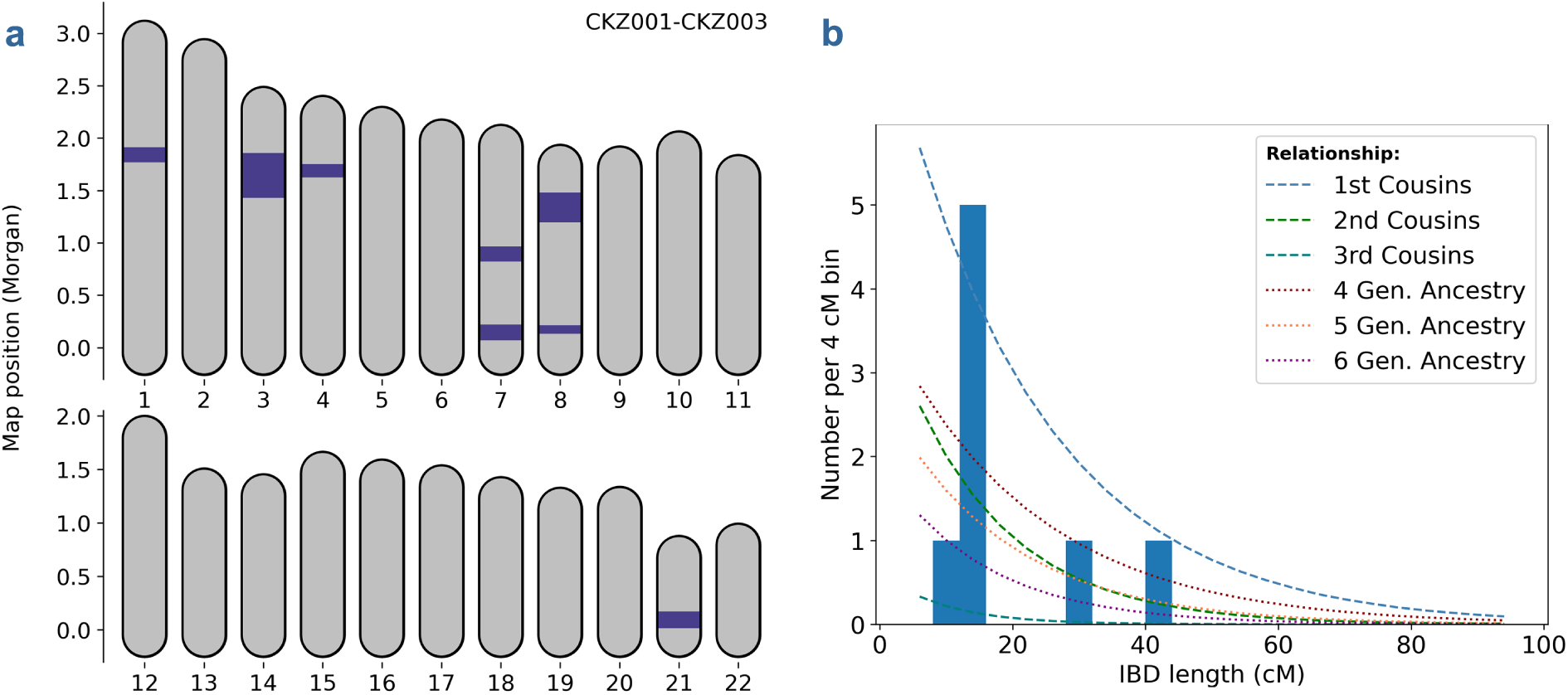
Shared IBD segments identified in two closely related Kenggir individuals. **Notes:** IBD segments longer than 4 cM were used for the plot. **a.** The blue color marks the physical locations of IBD segments on autosomal chromosomes; **b.** the histogram depicts IBD segments, where the curved lines indicate the expected densities of various degrees of kinship.

### Medieval IBD Network connects the Kazak Steppe and the Mongolian Plateau

After detecting genetic affinity between the Kenggir individuals and medieval Mongolians through PCA, outgroup *f_3_*-statistic, and Y-chromosome phylogenetics, we wanted to clarify whether the genetic relatedness was due to sharing a distant common ancestor or instead a recently shared genetic genealogy. To this end, we performed IBD (Identity by Descent) analysis with ancIBD, which can detect up to 6–8^th^ degree genetic relatedness^60^.

This analysis revealed that individuals from the Kazak steppe and the Mongolian plateau were connected through a more recent genetic genealogy, which resulted in at least one 8 cM IBD segment among the pairs analyzed. The individual pairs CKZ003-DA28, DA28-MRI001, CKZ002-CKZ003, and CKZ002-SHG003 shared 2–3 IBD segments, each at least 10 cM on average, indicating close genealogical relatedness. Relatives up to 6^th^ degree usually share two or more long IBD segments (>10cM)^61^.

Three individuals, CKZ003, DA28, and SHG003 had shared IBD segments with multiple individuals, implying the possibility of being ancestral to the other individuals analyzed (Fig 8 and Table S1). In the Y-chromosome phylogenetic tree (Fig 5b), SHG003 formed a clade together with the Kenggir individuals, whereas SHA001 clustered together with DA28 and KLK002. SHG003 shared two IBD segments with CKZ002, and one with CKZ003

**Fig 8.**
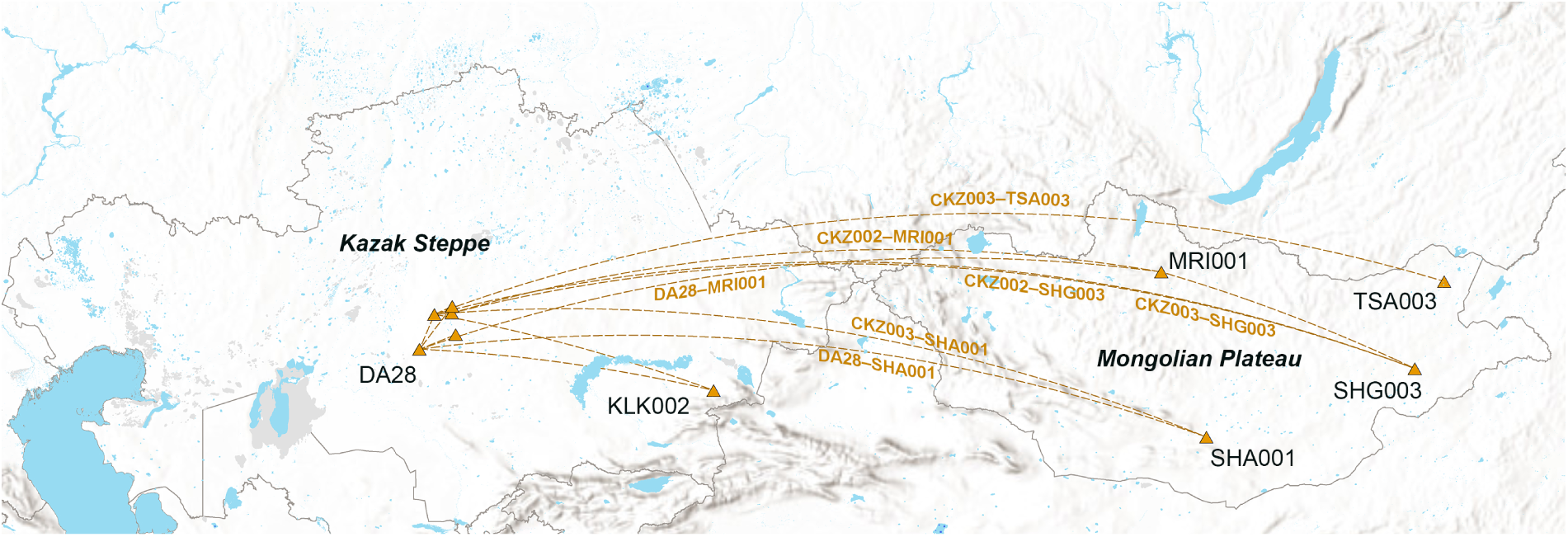
Medieval IBD Network connecting the Kazak Steppe and the Mongolian Plateau. **Notes:** The network was built among individual pairs who shared at least one IBD segment that is 8 cM or longer. Kenggir sites were not labeled, and the Kenggir IBD network is given in Fig S6.

As expected, the archaeologically younger individual (CKZ001) did not share any detectable IBD segments with SHG003 or any other medieval Mongolians. Both CKZ002 and SHG003 shared IBD segments with MRI001, which formed a distinct branch in the Y-chromosome phylogenetic tree. The female (CKZ004) did not share any detectable IBD segments with the Kenggir males but shared one segment with DA28. In light of these data, this female might have joined the Kenggir population as a bride.

One intriguing observation about the IBD network is that the medieval Mongolians apparently derived from a wide geographic area, encompassing North, South, and East of Mongolia. While they had limited connections among themselves, all of them showed connections with the Kenggir individuals and DA28 from the Ulitau massif. This observation could be explained in several ways: (1) before launching a military campaign westward to the Kipchak steppe, Genghis Khan unified the Mongolian Plateau and thus promoted admixture; (2) the westward campaign included soldiers and their households, including females and children, of various geographic origins; or (3) while the males were at the frontline, their family members were at the rear.

While the shared IBD segments between SHG003 and the Kenggir individuals (CKZ002 and CKZ003) could be resulted from shared paternal ancestry, all other IBD connections were most likely due to maternal relatedness through exogamy. Hundreds of years could separate Clade 1 and Clade 3 of the Y-chromosome tree (Fig 5b), which might show distinct medieval clans that were being connected through intermarriages.

### Relatively High level ROH found in a Kenggir individual

Runs of Homozygosity (ROH) provides us with information about the parental relatedness of an individual and it has implications for culture and health. In modern human populations, the distribution of ROH levels is uneven, with large populations, such as East Asians, having fewer ROH segments, while populations that practice consanguineous marriages, such as Balochi and Druze in Pakistan, as well as small, isolated populations, such as Koryak and Chukchi in Siberia, having more ROH segments on average^62^.

Similarly, ancient populations representing various archaeological cultures worldwide exhibit varying degrees of ROH^63^. We performed ROH analysis on modern and ancient individuals from Central Eurasia using hapROH program^63^. Two Kenggir individuals had detectable ROH segments that were at least 4 cM in size. One of the individuals, CKZ001, had 9 segments with the combined size of 83.2 Mb (Fig 9 a–c), which is comparable with the ROH seen in Yakut (Figure 9c). Over 88% of the offspring of first cousin marriages have longer than 20 cM ROH segments with combined length of at least 50 cM^63^. The similar levels of ROH can be found in 20% offspring of 2^nd^ cousin marriages but occur in less than 1% of the children of 3^rd^ cousin unions^63^.

**Fig 9.**
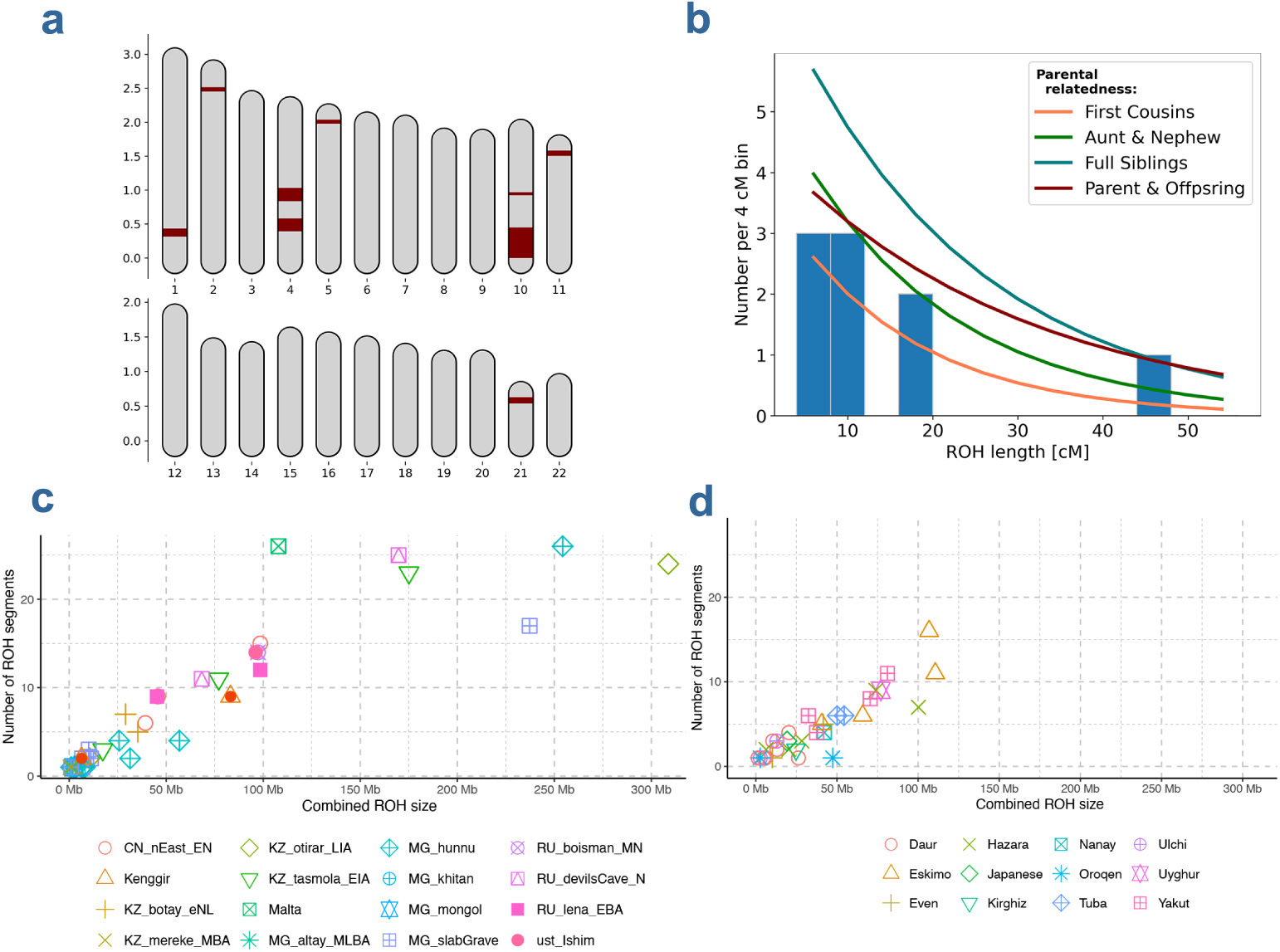
ROH identified in modern and archaeological individuals. **Notes: a** and **b** depict ROH segments found in CKZ001; **a.** the maroon-colored segments show physical positions of ROH on autosomal chromosomes; **b.** the ROH segments were depicted as a histogram, where the curved lines show the expected densities of various degrees of parental relatedness; Plot **c** depicts ROH in ancient individuals, while Plot **d** shows the ROH profiles found in modern individuals. The Kenggir individuals are marked with ochre dot in orange triangle.

At the early stages of the Golden Horde, individuals in a local population could trace their ancestry to various places in the Mongolian Plateau and other regions in Central Asia. At later stages, however, individuals could have shared more local ancestors and thus having higher chance of ROH. In addition, the strict exogamy tradition could become relaxed at least in some communities due to religious transition. Even with exogamy, however, if the mother and maternal grandmother of CKZ001 were related, then we would expect to see such levels of ROH.

### Genetic variants with anthropological significance

We analyzed several genetic variants in eleven medieval individuals, including the four Kenggir individuals and additional seven individuals from medieval period. Most of the individuals were connected via the IBD network (Fig 8). Frequencies of the genetic variants given in Fig 10 vary in human populations and their frequencies might be associated with cultural or environmental adaptation.

**Fig 10.**
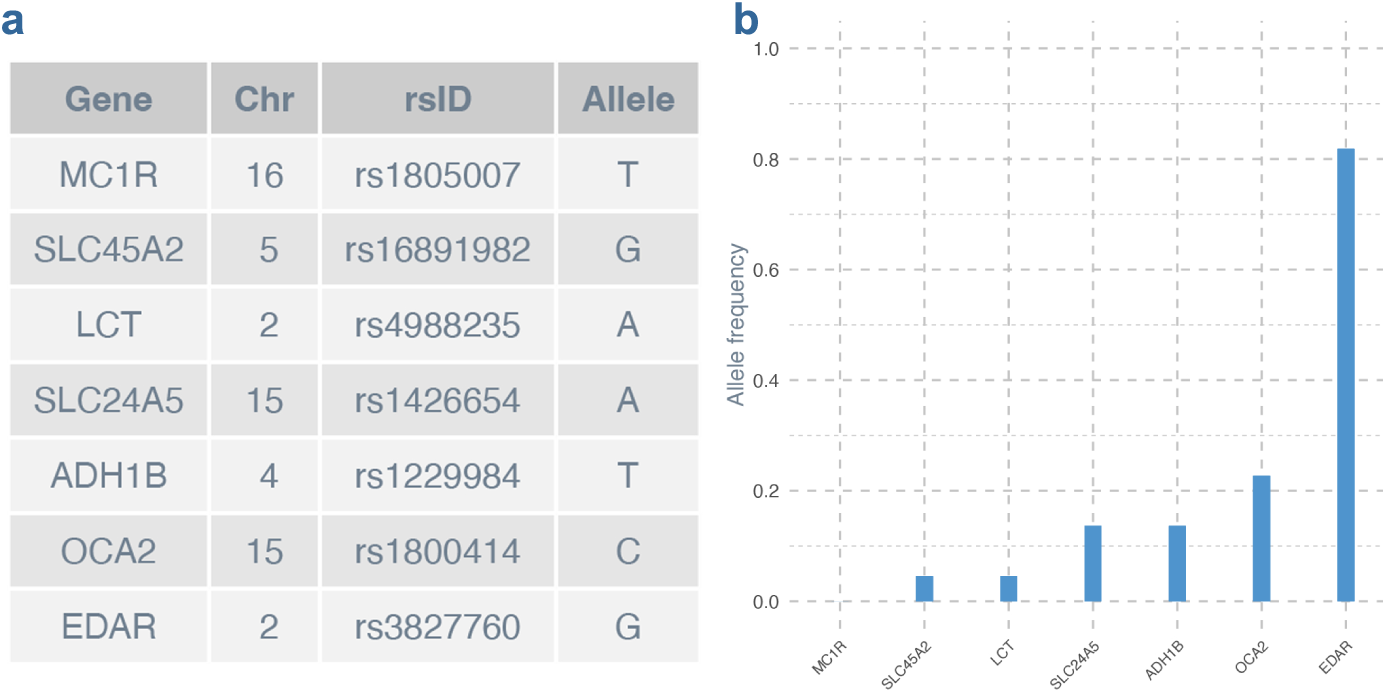
Genetic variants and their frequencies. **Notes:** The allele frequencies were calculated based on imputed ancient genomes of 11 individuals from Kazakstan and Mongolia.

As an example, the *LCT* allele A (rs4988235) has been hypothesized to be responsible for lactase persistence and occurs at high frequencies in Europeans, who consume milk as adults. The Kenggir individuals did not have this allele. Only one individual from Mongolia had heterozygous genotype. An alternative mechanism, such as gut microbiome composition, might be responsible for lactose digestion in the nomads of Central Eurasia^20^.

We also analyzed the alleles of four genes *MC1R, OCA2, SLC24A5,* and *SLC45A2* that are known to be associated with variation in human skin color pigmentation^64^. The *OCA2* allele C is found at high frequencies in East Asians and believed to be responsible for the light skin color of East Asians, suggesting that Europeans and Asians have developed light skin color through convergent evolution^65^. This specific *OCA2* allele does not occur in Europeans, although found at minuscular frequencies in Africans. In the medieval individuals analyzed, its frequency was just over 22%. Among the Kenggir individuals, only CKZ002 had this allele in heterozygous form.

The *SLC24A5* allele A (rs1426654) occurs at very high frequencies in Europeans, and it is also found in India^66^. Its frequency in the medieval individuals was 13.6%.

The frequency of the *MC1R* allele T in Europeans is 7.16%, but its frequencies in Asians and Africans are less than 0.5%. The medieval individuals did not have this derived allele.

The frequency of *SLC45A2* allele G in Europeans is 93.84%, but 0.6% in East Asians^64^, and its frequency in the medieval individuals was slightly higher than the frequency found in East Asians. Only one individual, CKZ004, was heterozygous for the allele.

The *EDAR* allele G (rs3827760) is responsible for hair-thickness^67^ and shovel-shaped incisors^68^ in East Asians and Native Americans. All medieval individuals had this allele, in either heterozygous or homozygous form. However, the frequency of *ADH1B* allele T in the medieval individuals was much lower than East Asian frequencies. The T allele of A*DH1B* is associated with a faster conversion of alcohol to acetaldehyde, resulting in facial flushing.

## Discussion

We leveraged historical, archaeological, and genetic data to confirm the Golden Horde affiliation of the four archaeological individuals and provided aDNA support for the association of Y-chromosome C3* (C2a1a3-F1918) haplotype with the Mongol Empire.

The whole genome analyses of the Kenggir individuals indicate that they mainly had Ancient Northeast Asian (ANA) ancestry, which is represented by the Neolithic individuals from the Devil’s Cave, northeast Asia^37^. When compared to modern human populations, the Kenggir individuals are more closely related with Altaic speaking (Mongolic, Tungusic, and Turkic) ethnic groups in north and northeast Asia. In addition, their genetic ancestry was supported by Y-chromosome haplotype analysis and the medieval IBD network that connects the Kazak steppe and the Mongolian plateau. As shown in the phylogenetic tree of Y-chromosome (Fig 5b), the Kenggir males share a common paternal ancestry with modern Kazaks, Mongols, Hazara, and other ethnic groups in Central Eurasia. The male buried in the Mausoleum of Joshi (CKZ002) harbored the previously hypothesized signature haplotype of Genghis Khan and was a paternal relative of both CKZ001 (Alasha Khan) and CKZ003 (Ayakkamir).

The archaeological findings showcase a population experiencing a cultural and religious transition from the traditional belief system rooted in Tängir to Islam. Constructing a mausoleum and placing the human remains in an east–west orientation, with the head toward the west in the direction of Mecca, aligns with the Islamic culture of Central Asia. However, interring animal heads and bones, saddles, clothes, and other items together with the deceased is non-Islamic. This burial practice of Kenggir individuals reflects a cultural transition, blending of traditional cultural elements with those of Islam. The gradual cultural and religious transition, which usually occurs side by side with linguistic shift, preceded the changes in the genomic make-up of individuals in the Kenggir population.

Even with whole genome data at hand, we could not reach an unambiguous conclusion with regard to the true identity of the male buried in the Mausoleum of Joshi. This will be an unattainable goal unless genetic data become available from the 12-13^th^ century pedigree of Genghis Khan. Populations or individuals who claim descent from Genghis Khan had various paternal lineages^17,42,44,45^, and no consensus has been reached.

From a historical perspective, based on the burial customs of the Mongol Empire, the Altan Urug elites were buried secretly without any monuments or mausoleums. Joshi might not be an exception to the imperial burial custom, given the fact that he passed away before his father, Genghis Khan deceased^69^. The Mausoleum of Joshi could have been built by a descendant of Joshi who believed in Islam, and dedicated and named the Mausoleum after Joshi Khan. Such a custom did exist in the Muslim communities in Central Asia^6^. Radiocarbon dates suggest a later time period than the lifetime of Joshi. As a recent study reported^31^, radiocarbon age estimates of the mausoleum’s building materials appeared younger than the lifetime of Joshi, as well.

From historical annals, we know that members of historical clans such as Kerey (Kereit), Jalayir, Dulat (Dughlat), and others served in the Mongol Empire and its successor states. For instance, the Jalayir Mukhali was a famous general of Genghis Khan in the 13^th^ century^8^, and another Jalayir, Kadir Ali Bek (*Қадырғали* in Kazak), served in the Kazak Khanate in the 16^th^ century^70^. In a different time and region, the Dughlat (Dulat) Mirza Haidar served as a general and governor in Moghulistan, a Central Asian successor state of the Mongol Empire that was ruled by the descendants of Chagatay, the second son of Genghis Khan^4^. A Yuan Dynasty mausoleum from northern China might belong to a nobleman of Turkic descent, as evidenced by the tombstone inscription, historical records, and his Y-chromosome haplogroup Q^19^.

These examples indicate that not only the direct descendants of Genghis Khan, but other figures were also part of the ruling elite of the Mongol Empire. Therefore, we cannot rule out the possibility that the Kenggir individuals might belong to a lineage other than Altan Urug. Future anthropological genetic and aDNA studies may help resolve the ambiguities around the identity of the individual interred in the Mausoleum of Joshi and the paternal lineage of Genghis Khan and his descendants.

Even though we cannot unequivocally reveal the identity of the Kenggir individuals, our data strongly suggest that they trace their ancestry to the Mongolian Plateau. The shared genomic profile of the Kenggir individuals and those from medieval Mongolia may reflect the genetic background of the ruling elites of the Mongol Empire, and quite possibly, that of Genghis Khan.

## Methods

### Archaeological sites and samples

For the archaeological excavation and aDNA analysis of the Golden Horde individuals from central Kazakstan, permissions were obtained from the Center for Protection of Historical and Cultural Relics, the Department of Culture, Archives and Documentation of Karaghandi, Kazakstan. Four archaeological individuals, three males and one female, were excavated by an archaeological team led by Dr. Aibar Kassenali. While two of the sites, Bolghan Ana and Ayakkamir, were excavated for the first time, the other two sites, Joshi and Alasha Khan, were excavated in the Soviet era, and the human remains were re-buried. We excavated these two sites again and sampled the human remains (Fig 1). Further details of the archaeological sites are given in the Supplementary Materials.

### Radiocarbon dating

Radiocarbon dating of the samples was carried out by Beta Analytic, Miami, FL, USA. A tibia fragment (CKZ001), a collarbone fragment (CKZ002), and rib fragments (CKZ003 and CKZ004) were used for radiocarbon dating (Fig 1e).

### Sequencing and authentication

Ancient DNA extraction, library preparation, and sequencing were performed in Japan. ***aDNA extraction:*** aDNA extraction was performed employing the method previously described^71^, with some modifications. aDNAs were extracted from the tooth or bone samples of CKZ001, CKZ003, and CKZ004. As for CKZ002, due to low sample quality, several libraries were prepared from aDNA extracted from a petrous bone and a fibula fragment (Table S1). ***Sequencing***: The aDNA libraries were subjected to an initial low coverage sequencing through MiSeq (Illumina). Thereafter, 100-bp paired-end reads were generated via HiSeq 2000 (Illumina). PCR duplicates were removed using MarkDuplicates from Picard toolkit^72^ and PickingBases^73^. **Authentication:** Sequencing error rate was estimated using the method as previously described^73^. The first and last few bases of sequence reads that had high rates of post-mortem damages specific to ancient genomes were removed with bamUtil^74^. Further information on aDNA libraries and sequences is available in Table S1.

### Variant calling and data compilation

Pseudo-haploid genotypes of the 1240K capture array sites were called from BAM files via the pileupCaller function of SequenceTools^75^, using the parameters -Q 30 -q 30. The resulted eigenstrat format data of the four Kenggir individuals were merged with a dataset of 400 individuals from the AADR (Allen Ancient DNA Repository, v54.1)^76^. The compiled dataset contained 404 individuals, including 328 ancient and 76 modern samples of Eurasian ancestry (Table S2).

### Analysis of uniparental lineages

Y-chromosome phylogenetic trees were built and haplotypes were identified via pathPhynder^77^ with default parameters. For calling Y-chromosome haplotypes, the reference human genome hs37d5 was used. Phylogenetic trees were constructed using the ISOGG Y-chromosome tree version 2019-2020^49^ as a reference. mtGenomes in FASTA format were created with the samtools^78,79^ consensus function using VCF files as input. The mtDNA VCF files were generated with snpAD^80^. We performed BLAST^81,82^ search on GenBank^83^ and retrieved ∼160 mtGenomes that had the highest levels of sequence similarity with the mtGenomes of the Kenggir individuals. mtDNA phylogenetic trees were built with RAxML-NG^84^, after aligning mtGenomes via MAFFT^85,86^. Mitochondrial haplotypes were called via Haplogrep 3.0^87^ online using BWA^88^ aligned muliti-fasta datasets as input. Plots were made with ggtree^89^ and ggplot2^90^ in R^91^.

### Kinship analysis

Kinship analysis was initially carried out via READ^59^, that is designed for detecting relatedness up to the 2^nd^ degree among archaeological individuals with low coverage aDNA data. READ accepts the TPED/TFAM format files of PLINK^92,93^. We performed additional kinship analysis via ancIBD, details of which were given in the IBD analysis section.

### Genetic structure analysis

Our dataset (n=404) was merged with a Human Origins (HO) dataset^94^, containing 997 modern individuals of Eurasian ancestry (Table S2). For cleaning the data, sites with 0.3 or more missingness and with minor allele frequency of 0.01 or lower were removed. In addition, only the individuals with a missingness less than 0.45 were kept (--geno 0.3 --mind 0.45 --maf 0.01). This resulted in a reduction of total number of SNPs of the original dataset (n=404) to ∼ 400K SNPs. PCA was performed on the cleaned dataset via smartPCA^33^, by projecting 404 individuals on the PCA space calculated by the HO dataset containg 997 individuals.

### Calculation of *f*-statistics

We used qp3Pop and qpDstat programs^34^ to calculate *f_3_* and *f_4_*-statistics, and to identify the individuals and populations that showed high levels of genetic affinity with the Kenggir individuals (Fig 3 & 4). For the *f*-statistic analysis, we cleaned our initial dataset with the parameter set [--geno 0.05 --mind 0.46]. Information on the samples included in the dataset is available in Table S2. Outgroup *f_3_*-statistic was calculated via *qp3Pop*^34^, with parameters *inbreeding: YES* and *outgroupmode: NO*. *f_4_*-statistic was calculated via *qpDstat*^34^. Both *qp3Pop* and *qpDstat* are from the ADMIXTOOLS package^34^. The *f3-*statistic pie charts and the background map were made with ArcGIS online, licensed to the University of Wisconsin-Madison.

### Estimation of ancestral components

We estimated ancestral genetic components of the Kenggir individuals and six other medieval individuals, two from Kazakstan and four from Mongolia using the *qpAdm* program from the ADMIXTOOLS package^34,35^. Except for Ust-Ishim, we used the same set of reference populations as Jeong and colleagues^20^. To be specific, we had nine reference populations: (1) Mbuti; (2) Italy_HG, paleolithic hunter-gathers lived in Italy; (3) Sardinians; (4) Turkey_N, Neolithic individuals from Anatolia; (5) Natufians, a Mesolithic population from Levant; (6) Iran_N, Neolithic individuals from the Iranian plateau, (7) Atayals, aboriginals from Taiwan, who positioned themselves in the southernmost location relative to other East Asians in the PCA space (Fig 2); (8) Mixe, a Native American group from Mexico; and (9) Ust-Ishim, which is represented by just a single individual from Siberia, found at the confluence of the rivers of Esil (Ishim) and Ertis (Irtysh).

In most cases, admixture models with a p-value higher than 0.05 are accepted (Table S4). The background map and the pie charts were created with ArcGIS online, licensed to the University of Wisconsin-Madison.

### IBD analysis

For performing IBD analysis via ancIBD^60^, we imputed the ancient genomes via GLIMPSE2^95,96^, using the default parameters and the 1000 Genomes Project phase 3 reference panel, which was created by surveying thousands of genomes of diverse ancestry aligned to the human reference genome build hs37d5. Imputed genomes were further analyzed with ancIBD using its default parameters designed for 1240K SNPs. The figures (Fig 7 a & b) were plotted with ancIBD. The IBD network map (Fig 8) was made with ArcMap, licensed to the University of Wisconsin-Madison.

### ROH analysis

ROH analysis was performed on a dataset of 269 individuals via hapROH program^63^. We performed ROH analysis separately for modern and ancient individuals, since the latter have pseudo-haploid genotypes. The first two graphs (Fig 9 a & b) were plotted with hapROH. The R package ggplot2^90^ was used for plotting the other two graphs.

### Analysis of genetic variants

VariantAnnotation^97^ was employed for retrieving genotypes from eleven imputed ancient genomes in VCF format. dbSNP (build 144) package^98^ in R^91^ was used for matching and confirming SNP position and rsIDs. For allele frequency calculation, an in-house R script was used. The R package ggplot2^90^ was used for plotting.

## Supporting information

Description of archaeological sites and it also includes supplementary figures.

Information on ancient DNA libraries and sequences.

Samples included in the datasets for PCA, f-statistic, and qpAdm analyses.

Ancestral components obtained via qpAdm analysis.

Information on the SNP markers used for placing the Kenggir individuals on the Y-chromosome phylogenetic tree.

IBD segments shared between the pairs of medieval individuals.

Information on the ROH segments found in the ancient and modern individuals from Central Eurasia.

## Data availability

BAM files of the ancient genomes will be made available upon publication of the manuscript in a journal.

## Acknowledgements

We are grateful to the Ministry of Culture and Sports, Kazakstan for supporting the project initiative. We appreciate the Center for Protection of Historical and Cultural Relics, the Department of Culture, Archives and Documentation of Karaghandi, Kazakstan for providing approval and assisting with the archaeological fieldwork. We are thankful to Tatiana Krupa for the reconstruction of the archaeological clothing and Aidos Esmagambetov for performing the craniofacial reconstruction of Bolghan Ana. The project was supported by a grant from the local government of Karaghandi oblast, awarded to A.K.; In addition, the project received support from a CRP grant (091019CRP2119) awarded to U.S. at Nazarbayev University and a grant awarded to S.N. at the National Institute of Genetics, Japan. Finally, we thank Theodore G. Schurr and Aaron Ragsdale for reviewing the manuscript and providing valuable comments.

## Author contributions

AA: Project design and coordination; data analysis; manuscript writing.

HK: aDNA extraction and library prep; aDNA sequencing and quality control; generation of final BAM files.

TK: aDNA extraction.

AK: Project design; archaeological excavation. SE: Archaeological excavation.

US: Reviewing the manuscript and providing input from a historical perspective. SS: Reviewing the manuscript and providing input from a population genetics perspective.

JH: Reviewing the manuscript and providing input from an anthropological perspective. NS: Project design; reviewing the manuscript and approving the final work for publication.

## Supplementary information

**Supplementary materials**: Description of archaeological sites and it also includes supplementary figures.

**Table S1:** Information on ancient DNA libraries and sequences.

**Table S2:** Samples included in the datasets for PCA, *f*-statistic, and qpAdm analyses.

**Table S3:** Ancestral components obtained via qpAdm analysis.

**Table S4:** Information on the SNP markers used for placing the Kenggir individuals on the Y-chromosome phylogenetic tree.

**Table S5:** IBD segments shared between the pairs of medieval individuals.

**Table S6:** Information on the ROH segments found in the ancient and modern individuals from Central Eurasia.

