## Supplementary material for "Genomes of the Golden Horde Elites and their Implications for the Rulers of the Mongol Empire": Description of archaeological sites and it also includes supplementary figures.

### Supplementary Materials

### Description of the archaeological sites

##### *Mausoleum of Alasha Khan*

We re-excavated the tombs in the mausoleums of Alasha Khan and Joshi Khan in 2018. Previously, these two archaeological sites were excavated by the archaeologist Margulan under the framework of the *Central Kazakhstan Archaeological Expedition* in 1947. In the first archaeological excavation, the tomb of a male (Fig S1a) and about 5 Kg of azure-colored glass materials were found in the Mausoleum of Alasha Khan[1]. Based on physical anthropological features, the male from the Mausoleum of Alasha Khan was estimated to be about 55 years old[2].

**Fig S1 | Schematic map of the Mausoleum of Alasha Khan and the Mausoleum of Joshi Khan**

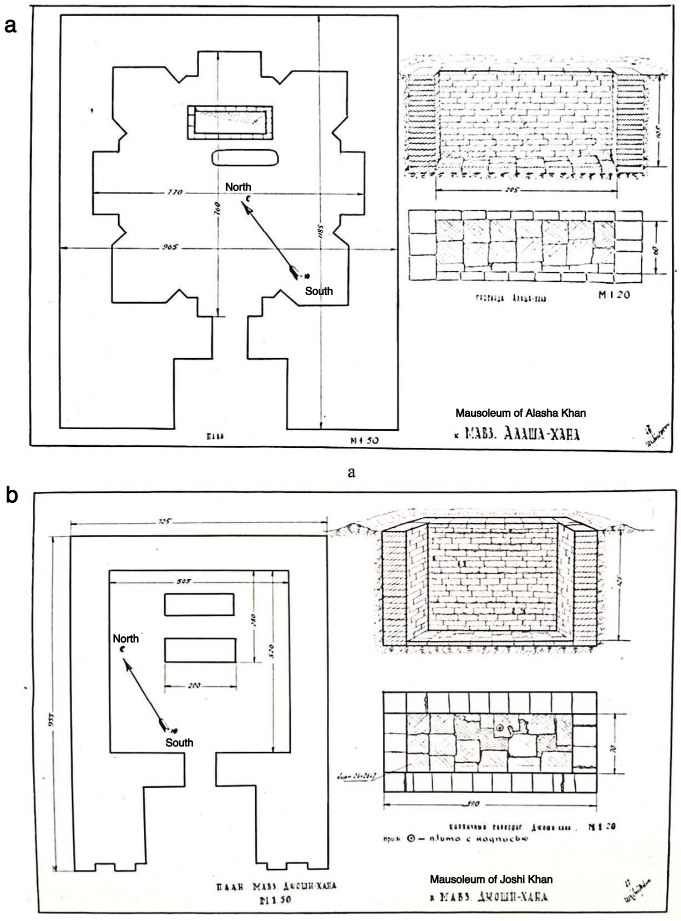


**Notes: a.** Mausoleum of Alasha Khan (CKZ001); **b.** The Mausoleum of Joshi (CKZ002). The unit of the measurements is cm. The figure was modified from Margulan (1947)[1].

##### *Mausoleum of Joshi Khan*

The expedition of Margulan found the remains of a male and a female in separate tombs located side by side in the Mausoleum of Joshi Khan (Fig S1b). The tombs were built of bricks, and the human bodies were placed in wooden coffins. In addition to human bones, animal bones, a camel skull, and remnants of leather and fabrics were discovered.
The male from the Mausoleum of Joshi Khan was estimated to be 40–45 years old, based on his skeletal features[2].

##### *Mausoleum of Ayakkamir*

The tomb inside the Mausoleum of Ayakkamir is rectangular, measuring 2×0.8 m, with a depth of 1.2 m. The edges and the top of the tomb were made of bricks, with its top being semi-dome shaped. Human skeleton, with no sign of disturbance, was found in the tomb. The body was oriented to the northwest, his arms were stretched down, and his legs were crossed. However, the skull was missing. A bronze cup was found lying near the right thigh. Among the soil piled on the surface of the tomb, coins and a small iron jar were found[3,4]. The discovery of coins indicates that the mausoleum served as a sacred place where many people came to worship prior to the establishment of the Soviet socialist system in Kazakhstan.

***Mausoleum of Bolghan Ana***

For the first time, comprehensive research was conducted at the Mausoleum of Bolgan Ana and unique archaeological artifacts were discovered. In the tomb, a female was lying supine, with her head facing northwest. She was in a robe, wearing a traditional Mongolian headware *boghta (богтаг)* and a pair of leather boots.

A metal mirror covered with leather and a saber were placed near her hip, and there was a wooden saddle near her feet (Fig S2). Among the artifacts found was a golden cup placed just above the right shoulder. There were also a hair stick and a pair of golden earrings with embedded white stone. The golden cup is round in shape, with protruding sides and a flat base. The diameter of the rim is 12 cm, and the height is 3.5 cm. The cup is made of an alloy of gold and copper. On the base of the inner surface, there is a flower motif, 6.5 cm in diameter. The golden earrings, made of thin gold wires, are 1.5 cm long, with a hook diameter of 1.7-1.8 cm. The hair stick, 8.2 cm long and 0.6 cm thick, is made from a bone of a small animal, with one end thickened and the other end pointed. The saber, 45 cm long and 0.5 cm thick, was forged from iron, with curved blade. The mirror is made of iron. Its diameter is 10 cm, its thickness is 0.8 cm, and its handle is about 12 cm long. It is heavily rusted and has broken into several pieces. One side was polished and used as a mirror[3,4].

**Fig S2 | Artifacts found in the tomb of Bolghan Ana**

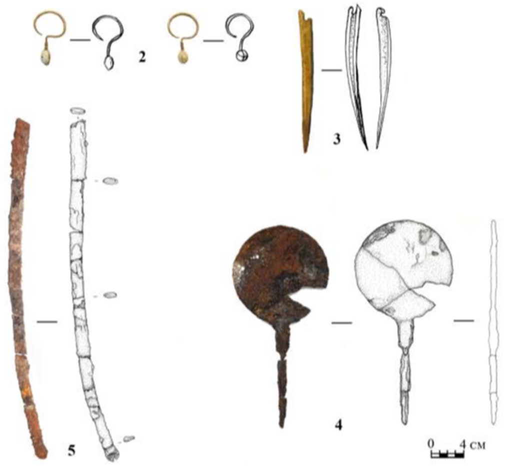

**Notes:** 2. Gold earrings; 3. Hair stick; 4. Metal mirror; 5. Saber

**Fig S3 | Fragments of the saddle found in the tomb of Bolghan Ana**

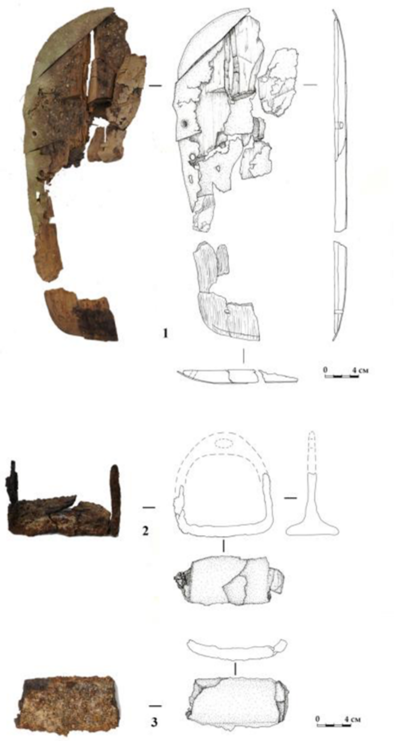

**Notes:** 1. Wooden frame of the saddle; 2 & 3. Stirrup

**Fig S4 | Kinship among the Four Archaeological Individuals**

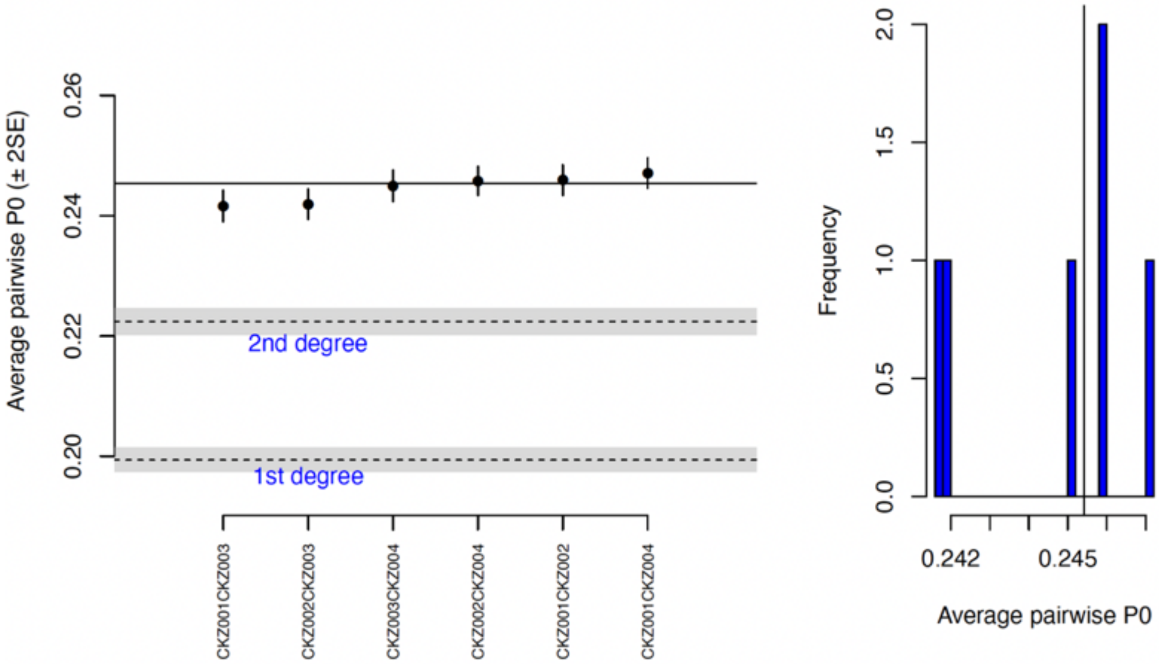

#### Fig S5 | IBD segments found in the pairs of CKZ001-CKZ002 and CKZ002-CKZ003

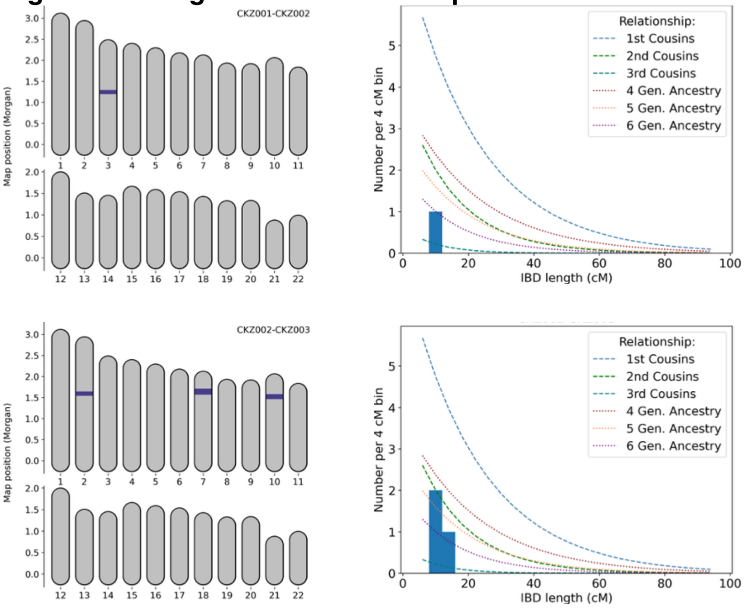


**Notes:** In the pairs, CKZ002 and CKZ003 are closer to each other. However, it is difficult to determine their actual relationship. CKZ001 and CKZ002 shared just a single IBD segment, indicating they might have belonged to the same population.

#### Fig S6 | IBD network among the medieval individuals from the Kazak steppe

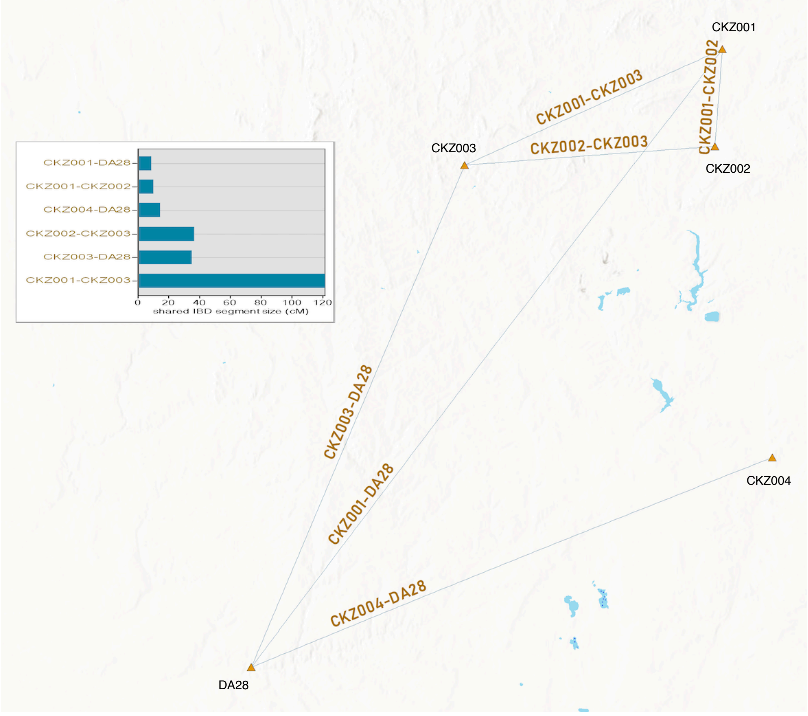


**Notes:** The bar plot indicates the total size of IBD segments shared by the individual pairs. The combined total size of IBD segments shared by CKZ001 and CKZ003 is ~ 120 cM. The IBD network was made via ArMap software licensed to UW-Madison.
